# Best Prediction of the Additive Genomic Variance in Random-Effects Models

**DOI:** 10.1101/282343

**Authors:** Nicholas Schreck, Hans-Peter Piepho, Martin Schlather

**Author notes:** Corresponding author: Chair of Stochastics and Its Applications, University of Mannheim, B6, 26, 68159 Mannheim, Germany.

## Abstract

The additive genomic variance in linear models with random marker effects can be defined as a random variable that is in accordance with classical quantitative genetics theory. Common approaches to estimate the genomic variance in random-effects linear models based on genomic marker data can be regarded as the unconditional (or prior) expectation of this random additive genomic variance, and result in a negligence of the contribution of linkage disequilibrium.

We introduce a novel best prediction (BP) approach for the additive genomic variance in both the current and the base population in the framework of genomic prediction using the gBLUP-method. The resulting best predictor is the conditional (or posterior) expectation of the additive genomic variance when using the additional information given by the phenotypic data, and is structurally in accordance with the genomic equivalent of the classical additive genetic variance in random-effects models. In particular, the best predictor includes the contribution of (marker) linkage disequilibrium to the additive genomic variance and eliminates the missing contribution of LD that is caused by the assumptions of statistical frameworks such as the random-effects model. We derive an empirical best predictor (eBP) and compare its performance with common approaches to estimate the additive genomic variance in random-effects models on commonly used genomic datasets.

## Introduction

The additive genetic variance is defined as the variance of the breeding value (BV) and is the most important determinant of the response of a population to selection (Falconer and Mackay 1996). The additive variance can be estimated from observations made on the population and is a principal component of the (narrow-sense) heritability, which is one of the main quantities of interest in many genetic studies (Falconer and Mackay 1996). The heritability is eminent, amongst other things, for the prediction of the response to selection in the breeder’s equation (Piepho and Moehring 2007; Hill 2010). Although non-additive genetic variation exists, most of the genetic variation is additive, such that it is usually sufficient to investigate the additive genetic variance (Hill *et al.* 2008).

More specifically, epistasis is only important on the gene level but not for the genetic variance (Hill *et al.* 2008), and Zhu *et al.* (2015) show that for human complex traits, dominance variation contributes little. Nevertheless, linkage disequilibrium (LD) is an important factor especially when departing from random mating and Hardy-Weinberg equilibrium, which is often the case in animal breeding (Hill *et al.* 2008; Dempfle 2018).

The additive genomic variance is defined as the variance of a trait that can be explained by a linear regression on a set of markers (de los Campos *et al.* 2015). Many authors have been chasing what is sometimes coined “missing heritability” (Maher 2008) which means that only a fraction of the “true” genetic variance can be captured by regression on influential markers. Initially, researchers have used genome-wide association studies (GWAS) in order to find quantitative trait loci (QTL) by single-marker fixed effect regression combined with variable selection. After having added the estimated corresponding genomic variances of the single statistically significant loci, they asserted that they could only account for a fraction of the “true” genetic variance. For instance, Maher (2008) found that only 5% instead of the widely accepted heritability estimate of 80% of human height could be explained. Golan *et al.* (2014) state that the “true” genetic variance is generally underestimated when applying variable selection, e.g. GWAS, to genomic datasets which are typically characterized by their high dimensionality, where the number of variables (markers) p is much larger than the number of observations n. It is well known that a lot of traits are influenced by many genes and that at least some loci with tiny effects are missed when using variable selection or even single-marker regression models. Consequently, Bernardo (1994) decided to fit all (RFLP-) markers in maize jointly using genomic best linear unbiased prediction (gBLUP), where he assumes the marker effect vector to be random. In animal breeding, Meuwissen *et al.* (2001) used Bayesian approaches (BayesA and BayesB) to fit all markers jointly in order to predict breeding values. Then, Yang *et al.* (2010) estimated the genomic variance in an approach that they termed genome-wide complex trait analysis genomic restricted maximum likelihood (GCTA-GREML) (Yang *et al.* 2011). They showed that quantifying the combined effect of all single-nucleotide polymorphisms (SNPs) explains a larger part of the heritability than only using certain variants quantified by GWAS methods. They illustrate their results on the dataset on human height by pointing out that they could explain a heritability, also termed “chip heritability” (Zhou *et al.* 2013), of about 45%. They concluded that the main reason for the remaining missing heritability was incomplete LD of causal variants with the genotyped SNPs, which refers to the general difference between the genetic variance and the genomic variance (Powell *et al.* 2010; de los Campos *et al.* 2015). However, the GCTA-GREML approach can be biased upwards as well as downwards (Wolc *et al.* 2013; de los Campos *et al.* 2015; Lehermeier *et al.* 2017; Fernando *et al.* 2017a). Recently, there has been a general discussion whether estimators for the genomic variance account for linkage disequilibrium (LD) between markers, which is defined as the covariance between the marker genotypes (Bulmer 1971). Some authors argue that estimators similar to GCTA-GREML lack the contribution of LD (Kumar *et al.* 2015, 2016; Lehermeier *et al.* 2017) whereas others (Yang *et al.* 2016) resolutely disagree. More specifically, Kumar *et al.* (2015, 2016) state that in GCTA-GREML the contributions of the p markers to the phenotypic values are assumed to be independent normally distributed random variables with equal variances. Thus, they claim that the random contribution made by each marker is not correlated with the random contributions made by any other marker which leads to a negligence of the contribution of LD to the additive genomic variance. In a study on the model plant Arabidopsis thaliana (The 1001 Genomes Consortium 2016), Lehermeier *et al.* (2017) use Bayesian ridge regression (BRR) to relate the phenotype flowering time to the genomic data. They use an estimator (termed M2) based on the posterior distribution of the marker effects obtained by Markov Chain Monte Carlo (MCMC) methods and show that this estimator explains a larger proportion of the phenotypic variance than the estimator, termed M1, based on gBLUP (VanRaden 2008; Yang *et al.* 2010, 2011). Lehermeier *et al.* (2017) argue that the reason for the better performance of the Bayesian estimator for the additive genomic variance (already mentioned in Sorensen *et al.* (2000); Zhou *et al.* (2013); Fernando and Garrick (2013); Fernando *et al.* (2017b)) is the explicit inclusion of linkage disequilibrium.

We show that the additive genomic variance in linear models with random marker effects (REM) can be defined as a random variable. Based on this premise, we propose a novel predictor of the additive genomic variance and place existing estimators in a joint framework permitting comparison with the new predictor. We contribute to the solution of many of the above mentioned controversies by reviewing common approaches to estimate the additive genomic variance, e.g. GCTA-GREML, and show that they estimate the unconditional (or prior) expectation of the random additive genomic variance. Combined with the assumptions on the unconditional distribution of the marker effects in the gBLUP-method this leads to an insufficient adaptation to the data and a negligence of the contribution of LD.

We introduce a novel best prediction approach for the additive genomic variance in both the current and the base population, i.e. we use the conditional (or posterior) expectation of the random additive genomic variance given the additional information by the phenotypic values for an improved adaptation to the data. We decompose the best predictor into the GCTA-GREML estimator and a function for the contribution of marker LD which determines whether GCTA-GREML is biased up- or downwards. The best predictor is structurally in accordance with the genomic equivalent of the additive genetic variance from classical quantitative genetics, i.e. it explicitly includes the contribution of LD. We propose an empirical best predictor (eBP) and illustrate our theoretical results on several commonly used genomic datasets.

## Material and Methods

### Linear Models

The connection of the *n*-vector *y* of phenotypic values and the mean-centered *n*-vector *g* of genomic values is given by

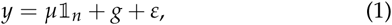

where *µ* denotes a fixed intercept, 𝟙_*n*_ := (1, …, 1) ^⊤^ is a *n*-row-vector containing 1’s, 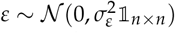 denotes environmental deviations, and 𝟙_*n×n*_ is the identity matrix of dimension *n*. For simplicity, we restrict the sample mean of the genotypic values to be 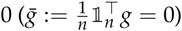.

In the following, we assume that the genome is mapped with *p* ∈ ℕ markers and we denote by **X** the *n* × *p* design matrix coding the genotypes of the markers. Then, the genomic values can be separated into the coded genotypes of the single markers and their corresponding *p*-vector *β* of marker effects:

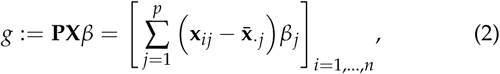

where 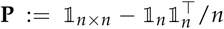 is the idempotent *n* × *n*-matrix used for column-wise mean-centering and **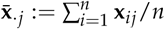** for *j* = 1, …, *p*. The restriction of the column-means of the marker genotype matrix to be 0 guarantees that the sample mean of the genomic values in (1) equals 0. This ensures the uniqueness of the definition of the vector *g* of genomic values in (2). Otherwise, different coding of the marker genotypes lead to different genomic values *g*.

Model (1) is called linear equivalent model (Henderson 1984) to the “standard” additive linear regression model

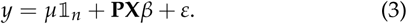

Model (3) allows for marker specific investigations and inferences on the genomic contribution to the phenotypic values, whereas estimation of parameters in model (1) has computational advantages.

Model (3) is a realization of *n* draws from the underlying data-generating process of the (mean-centered) marker genotypes (*X*_1_, …, *X*_*p*_) (Bühlmann and van de Geer 2011). This distribution as well as the corresponding genomic values in (2) relate to the current population of individuals.

When we are interested in the genomic values in the corresponding consistent base population, we should take the relationship (correlation) between the individuals into account (Powell *et al.* 2010; Legarra 2015). Assume that we have given a *n*× *n* relationship matrix **R**. Instead of the genomic values *g* or **PX***β* we investigate the uncorrelated genomic values defined by

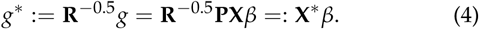

These are realizations of *n* draws from the underlying data-generating process 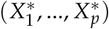 of marker genotypes in the base population. The sample mean of the genomic values in the base population, 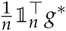, is usually different from 0.

The random-effects model is the statistical model that is probably most popular in genomic applications. Inferences on quantities based on genomic data in model (1) and (3) are often performed with the genomic best-linear-unbiased-prediction (gBLUP) method (Bernardo 1994). In this framework, we consider the single *p* components of the marker effect vector *β* in set-up (3) as independent normal random variables:

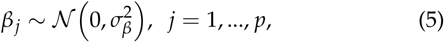

which implies that the effects are drawn at random from a common fixed normal distribution for each marker genotype.

In order to maintain the equivalence of models (1) and (3) we have to ensure the following equality in distribution:

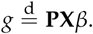

This can, for instance, be achieved by setting

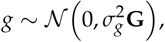

where 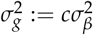 and

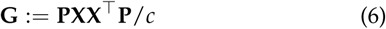

for some arbitrary *c >* 0. The *n* × *n*-matrix **G** is called genomic relationship matrix (GRM) and often *c* := 2 ∑ *p*_*j*_ (1 - *p*_*j*_), where *p*_*j*_ is the frequency of the minor allele at marker *j* (VanRaden 2008). For additional information on the gBLUP-method we refer to Appendix Genomic Best Linear Unbiased Prediction.

### Definitions of the Genomic Variance

We give an overview of the different approaches to define a genomic variance in the framework of the linear models (1) and (3).

Without further assumptions on the nature of the genome, we can define the sample variance

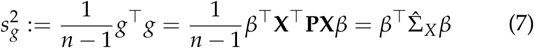

of the mean-centered *n*-vector of genomic values *g* = **PX***β*, see (2), in the current population (Ould Estaghvirou *et al.* 2013). Here, 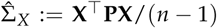 defines the sample variance-covariance matrix of the marker genotypes in the current population.

In the base population, we define the sample variance

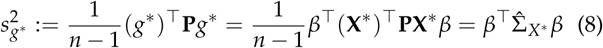

of the uncorrelated genomic values *g\** = **X***\*β*, see (4). Here, 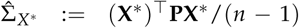 defines the sample variance-covariance matrix of the marker genotypes in the base population.

Alternatively, we can define the theoretical variance of the genomic values directly in the REM. The linear model is generated by drawing from the data-generating process of the marker genotypes (representative individual), and the model assumptions in the REM dictate that marker effects are random variables. This gives rise to three different sources of variance of the genomic values in the REM (marker genotypes random, marker effects random, or both random).

The additive genomic variance of a randomly sampled (representative) individual (Gianola *et al.* 2009; de los Campos *et al.* 2015; Fernando *et al.* 2017b) equals

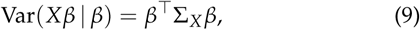

where Σ_*X*_ denotes the variance-covariance matrix of the marker genotypes.

The variance of a randomly sampled (representative) individual with random marker effects is given by

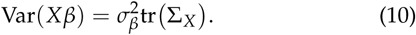

This is not the additive genomic variance (Gianola *et al.* 2009; de los Campos *et al.* 2015).

The variance of a randomly sampled trait averaged over individuals with fixed genotypes **X** equals

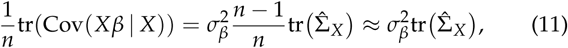

and does not equal the additive genomic variance.

We derive the equalities in (9), (10) and (11) in more detail in the Appendix Theoretical Variances of the Genomic Values in the REM. These quantities refer to the genotypes in the current population. We can apply the same definitions in the base population by considering the data-generating process of the genotypes in the base population (exchange *X* by *X\**).

In Table 1 we give an overview of the different possibilities to define the variance of the genomic values in the REM.

**Table 1.** Overview of different definitions of the variance of the genomic values in the current population and their expression in the random-effects model. Analogous quantities for the base population can be obtained by exchanging *X* by *X*\*. The sample variance 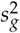 and the theoretical variance Var(*Xβ|β*) define the sample and theoretical version of the additive genomic variance.

| Variance of Genomic Values |  |  |  |
| --- | --- | --- | --- |
| Sample Variance | Theoretical Variance |  |  |
| $s_g^2$ | $\text{Var}(X\beta)$ | $\text{tr}(\text{Cov}(X\beta\vert X))$ | $\text{Var}(X\beta\vert \beta)$ |
| $\beta^\top \hat{\Sigma}_X \beta$ | $\sigma_\beta^2 \text{tr}(\Sigma_X)$ | $n\sigma_\beta^2 \text{tr}(\hat{\Sigma}_X)$ | $\beta^\top \Sigma_X \beta$ |

**Table 2.**
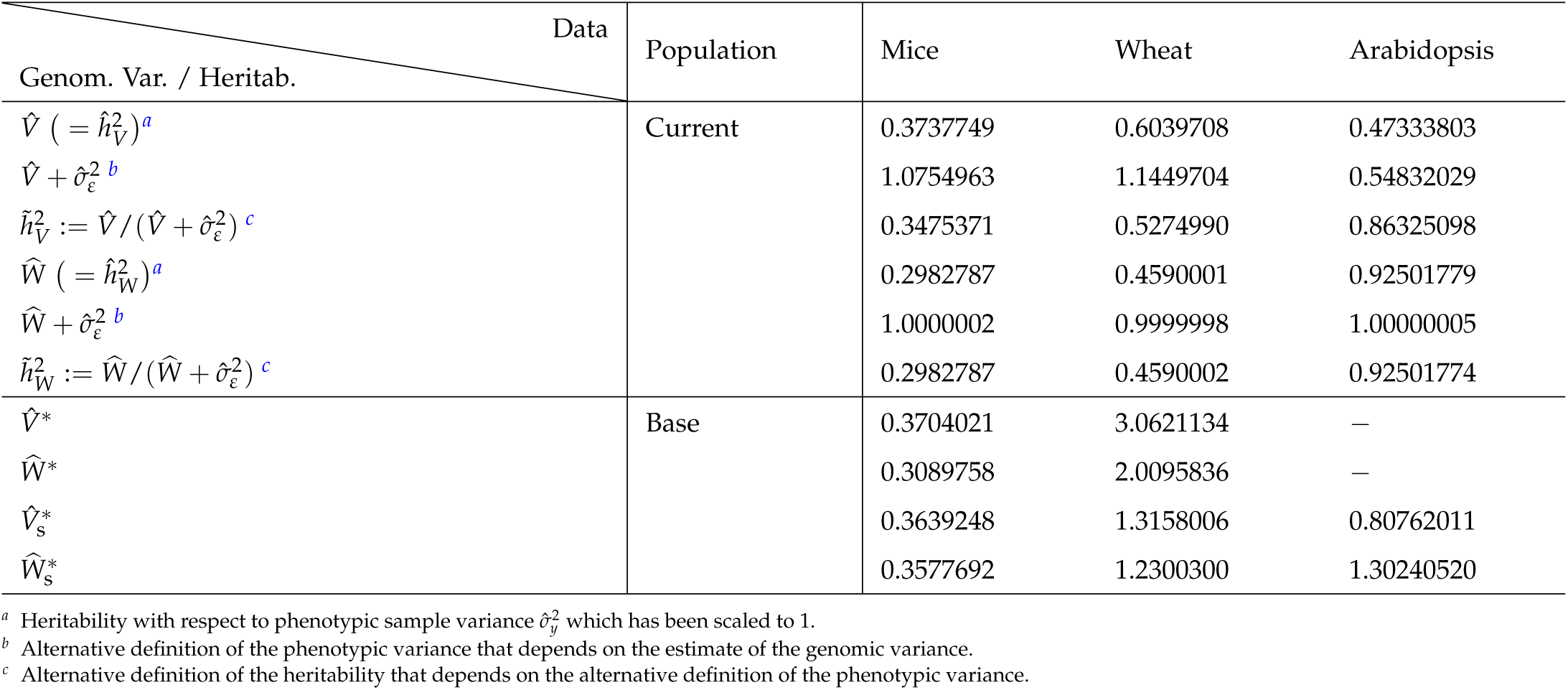
Estimation results for the unconditional expectation *V* and the best predictor *W* for the additive genomic variance in the current population for the mice, wheat, and Arabidopsis datasets. We also present the corresponding heritabilities with respect to the sample variance of the phenotypic values and with respect to the sum of the additive genomic and residual variance 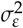. In addition, we depict the estimation results for the unconditional expectation *V*^*∗*^ (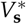 when using the GRM for the transformation) and the best predictor *W*^*∗*^ (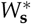 when using the GRM for the transformation) for the additive genomic variance in the base population.

In actual applications, we have to replace Σ_*X*_ in (9) by its estimator 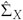. Consequently, the sample variance (7) as well as the theoretical (9) effectively represent the additive genomic variance, the genomic equivalent of the additive genetic variance (Bulmer 1971; Falconer and Mackay 1996), in the current population. In the following, we do not explicitly distinguish between the sample or the theoretical version of the variance, and will speak only of the additive genomic variance.

In the following, we focus on the estimation of the additive genomic variance in the general form

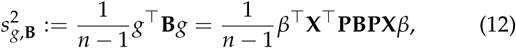

which is a non-negative quadratic form of the genomic values. By specifying the positive semi-definite and symmetric _*n*× *n*_-matrix **B** we determine whether the genomic variance refers to the current population (**B** = 𝟙_*n*× *n*_), see (7), or the base population (**B** = **R**^-0.5^**PR**^−0.5^), see (8). Because the randomness of the marker genotypes is not explicitly necessary to derive (12), we can easily express all results in the terminology of the genomic values *g* defined in the equivalent model.

In the framework of the REM, the marker effects *β* in model (3) and the genomic values *g* in model (1) are random variables. Consequently, the additive genomic variance in (12) is also a random variable, and has to be predicted in an optimal way before finally being estimated.

First, we will show that estimators for the unconditional expectation of (12), like GCTA-GREML, are of the form (10) and (11), and therefore do not estimate the additive genomic variance.

Then, we introduce the (frequentist) best predictor for the additive genomic variance 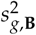 and show that this approach maintains the structure of the additive genomic variance in (12), the genomic equivalent of the additive genetic variance.

### The Expectation of the Additive Genomic Variance

The expectation of the random variable 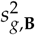 in (12) minimizes the quadratic form

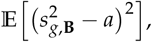

with respect to all real numbers *a*, i.e. 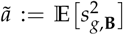 is the best approximation of 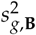 in the absence of additional information (van der Vaart 2007). The unconditional (or prior) expectation of 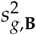 equals

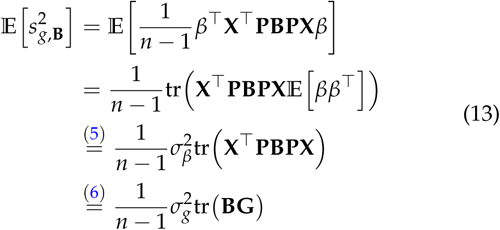

because of the properties of the trace.

For the additive genomic variance in the current population, 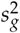, we choose **B** = 𝟙_*n×n*_ in (13) and obtain

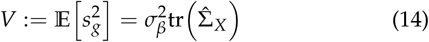

in model (3) or

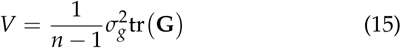

in the equivalent model (1). Unconditional expectations of the form *V* for the additive genomic variance are considered in Ould Estaghvirou *et al.* (2013), for example.

For the additive genomic variance in the base population, 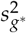*\**, we choose **B** = **R**^−0.5^ **PR**^−0.5^ in (13) and obtain

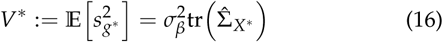

in model (3) or

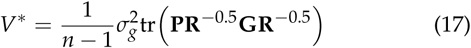

in the equivalent model. Often (VanRaden 2008; Yang *et al.* 2010, 2011; Speed *et al.* 2012; Vinkhuyzen *et al.* 2014; Legarra 2015), the matrix **R** used for the transformation to the base population is assumed to be the GRM **G** defined in (6). Then, the unconditional expectation *V*\* of the additive genomic variance simplifies to

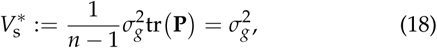

and the variance component 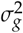 from the gBLUP-method is considered as the (unconditional expectation of the) additive genomic variance in the base population. We recommend caution when using this simplification because the GRM **G** is in general singular (because **P** is singular), and therefore **G**^−1^ is not well defined.

We emphasize that only the diagonal elements of the sample variance-covariance matrix (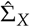 or 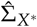) of the marker genotypes influence the unconditional expectations *V* and *V*\* of the additive genomic variance. The model assumptions in the REM dictate the matrix 𝔼[*ββ*^*T*^] to be diagonal which leads to a negligence of the off-diagonal elements of 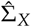 or 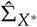 in (13). The covariances (LD) between the marker genotypes are not included and *V, V∗* and 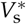 are of the same form as Var(*Xβ*) in (10) and 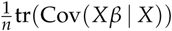 in (11). This implies that the unconditional expectation of the random additive genomic variance 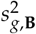 is structurally not fully in accordance with the additive genomic variance.

Explicit formulae for the estimation of the unconditional expectations *V, V*\* and 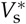 will be given in the Appendix Estimation of the Additive Genomic Variance in the REM.

### Best Prediction of the Additive Genomic Variance

The unconditional expectation of 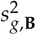 in (13) is strongly influenced by the model assumption on the marginal distribution of the marker effects and does not use additional information given by the phenotypic values *y* in model equations (1) and (3). By contrast, the conditional expectation, given the phenotypic values *y*, can make use of the information in *y*.

Generally, the conditional (or posterior) expectation of a random variable *Z* (in our case 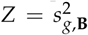) given the knowledge of the random vector *Y* is defined as the “best prediction” (Searle *et al.* 1992; van der Vaart 2007) of the random variable *Z*. The best predictor

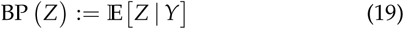

is the unique function *g*_0_(*Y*) that minimizes the mean square error of prediction

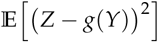

over all functions in *Y*, i.e. the conditional expectation is the projection (closest element in a given set of functions) of *Z* onto the linear space of all functions in *Y* (Searle *et al.* 1992; van der Vaart 2007).

The best predictor in (19) is by definition an unbiased predictor for the random variable *Z* and *g*_0_(*Y*) maximizes the correlation Cor(*Z, g*(*Y*)), i.e. we can replace the target random variable *Z* by the best predictor defined in (19) in an optimal way (Searle *et al.* 1992). Instead of inferring the unobservable target random variable, we conduct inferences on the best predictor. Because the best predictor has realized in a given dataset (*Y* = *y*), it is estimable (Searle *et al.* 1992).

In the following, we introduce a novel approach of considering the frequentist best predictor instead of the unconditional expectation for the random additive genomic variance 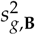 in (12). We proceed according to (19) and define

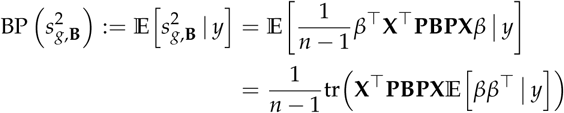

for the given phenotypic values *y*, because **X** is constant and therefore independent of *y*. The matrix of conditional second moments of the marker effects *β* is usually non-diagonal (contrary to 𝔼[ *ββ*]^*T*^) and can be expressed as

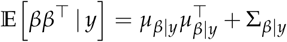

using the BLUP *µ*_*β|y*_ := 𝔼 [*β|y*] of the random vector *β* and the variance-covariance matrix Σ_*β y*_ := Cov *β y* of *β* given the data *y*. Then, the best predictor equals

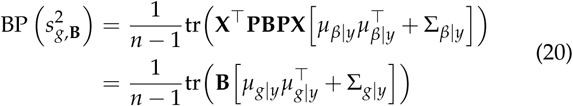

where the last equality holds because of the connection

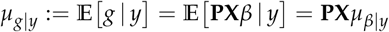

of the BLUPs and the conditional variance-covariance matrices

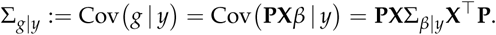

in models (1) and (3), see also Appendix Genomic Best Linear Unbiased Prediction.

For the best predictor of the additive genomic variance in the current population we set **B** = 𝟙_*n×n*_ in (20) and obtain

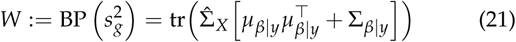

in model (3) or

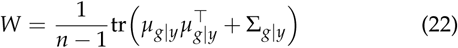

in the terminology of the equivalent model (1).

For the best predictor of the additive genomic variance in the base population we set **B** = **R**^−0.5^ **PR**^−0.5^ in (20) and obtain

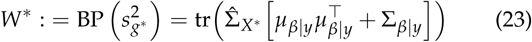

in model (3) or

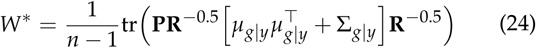

in the terminology of the equivalent model (1).

We emphasize that the best predictor of the additive genomic variance in the current population (*W*) as well as in the base population (*W*\*) includes the contribution of all elements of the sample variance-covariance matrix of marker genotypes (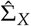 or 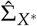), and hence comprise LD information, contrary to the unconditional expectations *V, V*\*, and 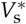 of the additive genomic variance from the previous section.

Explicit formulae for the empirical best predictors (eBP) of the additive genomic variance as well as a formula for 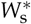 (approximate approach using the GRM **G** for transformation to the base population) will be given in the Appendix Estimation of the Additive Genomic Variance in the REM. We compare the use of the unconditional expectation and the best predictor for the prediction of the random additive genomic variance in the REM in Tables 3 and 4 in the Appendix.

**Table 3.** Overview of Prediction Approaches for the Random Additive Genomic Variance in the Random-Effects Model with the gBLUP-method 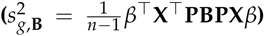. **X** is the matrix of marker genotypes, **P** the matrix for column-wise mean-centering, **B** a positive semi-definite matrix, 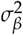 the variance component of the marker effects *β, μ*_*β|y*_ the BLUP of *β*, Σ_*β|y*_ the conditional covariance matrix of *β* given the phenotypic data *y*, **R** a relationship matrix, 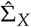 the sample variance-covariance matrix of the marker genotypes in the current population, 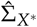 the sample variance-covariance matrix of the marker genotypes in the base population.

|  | Unconditional Expectation | Best Prediction |
| --- | --- | --- |
| **General Formula** | $\mathbb{E}[s_{g,B}^2] = \frac{1}{n-1} \sigma_\beta^2 \text{tr}(\mathbf{X}^\top \mathbf{P} \mathbf{B} \mathbf{P} \mathbf{X})$ | $\text{BP}(s_{g,B}^2) = \frac{1}{n-1} \text{tr}(\mathbf{X}^\top \mathbf{P} \mathbf{B} \mathbf{P} \mathbf{X} [\mu_{\beta\vert y} \mu_{\beta\vert y}^\top + \Sigma_{\beta\vert y}])$ |
| **Current population** | $V = \sigma_\beta^2 \text{tr}(\hat{\Sigma}_X)$ | $W = \text{tr}(\hat{\Sigma}_X [\mu_{\beta\vert y} \mu_{\beta\vert y}^\top + \Sigma_{\beta\vert y}])$ |
| **Base Population** | $V^* = \sigma_\beta^2 \text{tr}(\hat{\Sigma}_{X^*})$ | $W^* = \text{tr}(\hat{\Sigma}_{X^*} [\mu_{\beta\vert y} \mu_{\beta\vert y}^\top + \Sigma_{\beta\vert y}])$ |
| **Features** | • Best approximation of the additive genomic variance in the absence of information<br>• No inclusion of LD | • Best approximation of the additive genomic variance using additional information given by phenotypic values (adaptation to the data)<br>• Explicit inclusion of LD<br>• Orthogonal decomposition of the phenotypic variance in the current population (unique definition of the heritability)<br>• Genomic equivalent of the additive genetic variance |

**Table 4.** Overview of Prediction Approaches for the Random Additive Genomic Variance in the Random-Effects Model with the gBLUP-method in the equivalent version of the linear model 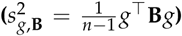. G is the genomic relationship matrix, **P** the matrix for column-wise mean-centering, **B** a positive semi-definite matrix, 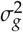 the variance component of the genomic values *g, μ*_*g|y*_ the BLUP of *g*, Σ_*g|y*_ the conditional covariance matrix of *g* given the phenotypic data *y*, **R** a relationship matrix.

|  | Unconditional Expectation | Best Prediction |
| --- | --- | --- |
| **General Formula** | $\mathbb{E}[s_{g,B}^2] = \frac{1}{n-1} \sigma_g^2 \text{tr}(\mathbf{B} \mathbf{G})$ | $\text{BP}(s_{g,B}^2) = \frac{1}{n-1} \text{tr}(\mathbf{B} [\mu_{g\vert y} \mu_{g\vert y}^\top + \Sigma_{g\vert y}])$ |
| **Current population** | $V = \frac{1}{n-1} \sigma_g^2 \text{tr}(\mathbf{G})$ | $W = \frac{1}{n-1} \text{tr}([\mu_{g\vert y} \mu_{g\vert y}^\top + \Sigma_{g\vert y}])$ |
| **Base Population** | $V^* = \frac{1}{n-1} \sigma_g^2 \text{tr}(\mathbf{P} \mathbf{R}^{-0.5} \mathbf{G} \mathbf{R}^{-0.5})$ | $W^* = \frac{1}{n-1} \text{tr}(\mathbf{P} \mathbf{R}^{-0.5} [\mu_{g\vert y} \mu_{g\vert y}^\top + \Sigma_{g\vert y}] \mathbf{R}^{-0.5})$ |
| **Features** | • Best approximation of the additive genomic variance in the absence of information<br>• No inclusion of LD<br>• Transformation with GRM: $\sigma_g^2$ replaces $V^*$ | • Best approximation of the additive genomic variance using additional information given by phenotypic values (adaptation to the data)<br>• Explicit inclusion of marker LD<br>• Orthogonal decomposition of the phenotypic variance in the current population (unique definition of the heritability)<br>• Genomic equivalent of the additive genetic variance |

### Statistical Analysis (Genomic Data)

For an illustration of the theoretical results of the previous sections we used the mice dataset that comes with the *R*-package BGLR (Perez and de los Campos 2014). The data originally stem from an experiment by Valdar *et al.* (2006a,b) in a mice population. The dataset contains the matrix **X** with values in {0, 1, 2 of *p* = 10346 polymorphic marker genotypes that were measured in *n* = 1814 mice. The trait (*n*-vector *y*) under consideration was body length (BL). The relationship of the mice is recorded in the *n × n* pedigree matrix **R** and is used for the transformation to the base population.

Additionally, we used the publicly available historical wheat dataset that also comes with the *R*-package “BGLR” (Perez and de los Campos 2014). The data originally stems from CIMMYT’s Global Wheat Program and consists of *n* = 599 lines of wheat where the trait under consideration was average grain yield. The phenotypes are divided up into four basic target sets of environments designated as Wheat I, Wheat II, Wheat III and Wheat IV where we only considered the first one. The dataset contains the matrix of marker genotypes for *p* = 1279 markers as well as a relationship matrix.

Moreover, we analyzed a population of *n* = 1057 fully sequenced Arabidopsis lines for which phenotypes and genotypes are publicly available by the effort of the Arabidopsis 1001 Genomes project (The 1001 Genomes Consortium 2016). The lines represent natural inbred lines and we examined the same trait, namely flowering time at 10°C (FT10), and the same *p* = 193697 SNP-markers that were used in Lehermeier *et al.* (2017). For these data no relationship matrix was available.

For each dataset, we used the gBLUP-method in the equivalent version (computational advantages) implemented in the *R*-package “sommer” (Covarrubias-Pazaran 2017) to fit a REM. We worked with the option REML (restricted maximum likelihood) to obtain estimates (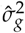 and 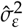) for the variance components. The method also returned estimates of the best predictor of the genomic effects *µ*_*g|y*_ and the their variance-covariance matrix 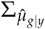.

We used this outcome for the estimation of the unconditional expecation *V* and the BP *W* of the additive genomic variance in the current population and the as well as the unconditional expectation *V*\* and the BP *W*\* for the additive genomic variance in the base population (except for the Arabidopsis dataset, where no relationship matrix was available). Although the GRM is not invertible, we will show in the Appendix Estimation of the Additive Genomic Variance in the REM how to use the GRM for a transformation to the base population, and to calculate the corresponding unconditional expectation 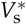 and the BP 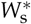 for the additive genomic variance in the base population.

We conducted all calculations with the free software R (R Development Core Team 2017). Detailed information about the calculations as well as the programming code with its output is provided in the supplemental File S1.

## Data Availability

The authors affirm that all data necessary for confirming the conclusions of this article are represented fully within the manuscript and the supplemental material that has been uploaded to figshare. Supplemental File S1 contains a detailed description of the estimation of the genomic variances for the gBLUP-method as well as the corresponding *R*-code and its output.

## Results

In the first section of Table 2 we present the estimation results for the unconditional expectation *V* and the best predictor *W* for the additive genomic variance in the current population. In the mice and wheat dataset 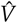 exceeds *Ŵ*, whereas for the Arabidopsis data, the empirical best predictor is about double the size of the unconditional expectation.

The sample variance of the phenotypic values has been scaled to 1. The sum of 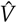 and the residual variance is larger than the phenotyic variance for the mice and wheat data but smaller for the Arabidopsis data. Technically, it is possible to define the heritability in two ways, namely with respect to the phenotypic variance and with respect to the sum of the additive genomic variance and the residual variance. The sum of the emprical best predictor *Ŵ* and the residual variance, however, equals the scaled phenotypic variance of 1 remarkably exactly for all datasets considered.

In the second section of Table 2 we first present the estimation results for the unconditional expectation *V*\* and the best predictor *W*\* for the additive genomic variance in the base population using the given relationship matrices for the transformation. For the mice data, 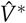 and *Ŵ*\* are similar to their analogons 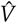 and *Ŵ* in the current population. For the wheat data, however, the estimated unconditional expectation and empirical best predictor in the base population are about five times larger than those in the current population and exceed the sample phenotypic variance in the current population. By this approach, it is not possible to define a heritbility in the base population because both the estimate of the residual variance and the phenotypic sample variance refer to the current population.

The estimation results for the unconditional expectation 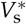 and the best predictor 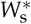 for the additive genomic variance in the base population using the GRM for the transformation differ from those using the given relationship matrices by a considerable amount. In the mice and wheat data, 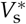 is larger than 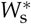, whereas for the Arabidopsis data the empirical best predictor exceeds the estimated unconditional expectation. This conforms to the behavior of *V* and *W* in the current population.

## Discussion

We have shown that commonly used estimators for the additive genomic variance in the REM with genomic marker data are based on the unconditional expectation of the random additive genomic variance. We have introduced a novel best prediction approach for the random additive genomic variance in both the current and the base population. In the following, we discuss several important implications.

### Current and Base population

Common ways of estimating the additive genomic variance focus on the base population. These approaches are independent of the actual current population and consequently valid even if the generations change.

If one aims at the response of a population to selection, however, it might be more meaningful to estimate the additive genomic variance in the actual given population. This implies that the estimation of the genomic variance has to be conducted again when the individuals change. A formal definition of the heritability is best possible in the current population, where the phenotypic and residual variance are estimable.

We have preferred to use given relationship matrices for the transformation of the genomic values to the base population. In the case that such a matrix is not available, we have shown how to use genomic relationship matrices for the transformation, although a formal inversion of GRM’s is in general not possible.

In Table 2 we have illustrated that we can decompose the sample phenotypic variance into the sum of the empirical best predictor of the additive genomic variance in the current population and into the estimated residual variance. This is due to the orthogonal projection property of the conditional expectation which gives the best approximation of the random additive genomic variance. This enables a unique definition of the heritability in the current population. It is never possible, however, to transfer the residual variance to the base population. Consequently, a definition of the heritability in the base population is not straight-forward.

### The gBLUP-method and the Bayesian approach

The frequentist gBLUP-method can also be set-up in the context of Bayesian regression models (with prior distribution for the effect vector as defined in (5) and uninformative priors for the variance components). Lehermeier *et al.* (2017) considered the additive genomic variance 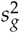 in the current population, see (7), and used Bayesian ridge regression to estimate

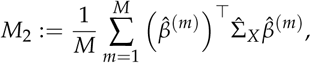

where 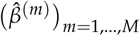 denotes MCMC samples from the posterior distribution of *β*. In that approach, Lehermeier *et al.* (2017) have estimated the posterior mean of the additive genomic variance 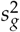 in the current population. This approach is the Bayesian equivalent of the (frequentist) empirical version of the best predictor of 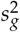 in (12) in the current population. *M*2 does not describe the genomic variance in the base population and should not directly be compared with approaches introduced e.g. in Yang *et al.* (2010, 2011). Analogously to the best predictor *W*^*∗*^, see (23), for the genomic variance in the base population, one can consider

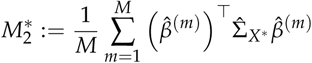

as the posterior mean of the genomic variance in the base population in Bayesian regression models.

The frequentist gBLUP-method provides a more formal approach to the prediction of the random additive genomic variance in linear models with random effects than the Bayesian approach. It enables the derivation of explicit formulas for the predictors (unconditional expectation and best predictor) of the random additive genomic variance using the standard output of the gBLUP-method which goes hand in hand with a fast implementation of the empirical version of the predictors. The connection between the BLUP *μ*_***β****|y*_ and its covariance for the random marker effects with the additive genomic variance are clearly visible. This enables us, for instance, to derive the decomposition of the best predictor of the random additive genomic variance into the unconditional expectation and a function for the marker LD in the following section.

### Influence of Linkage Disequilibrium

In Section Definitions of the Genomic Variance we have seen that the (random) additive genomic variance equals

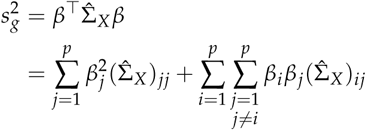

in the current population, and

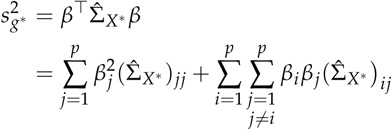

in the base population. We emphasize that the variance-covariance matrix of the marker genotypes (marker LD) plays a decisive part in the determination of the additive genomic variances in both the current and the base population. The variances 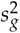 and 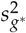 are structurally in accordance with the classical additive genetic variance (Bulmer 1971; Falconer and Mackay 1996), which is caused by the genotypes whereas the genotypic effects are fixed.

In the REM, however, the marker effects are random with unconditional expectation 0 and unconditional diagonal variance-covariance matrix with equal variances 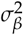. As a consequence, the unconditional expectation of the additive genomic variance

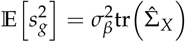

in the current population and

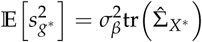

in the base population contain only the variances of the marker genotypes in the corresponding population. In addition, the unconditional expectation resembles both the variance of a randomly sampled trait for a randomly sampled individual and the variance of a randomly sampled trait for individual with fixed genotypes, see Table 1 for an overview.

We show in the Appendix Estimation of the Additive Genomic Variance in the REM that we can partition the best predictor of the random additive genomic variance 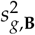 in the following way:

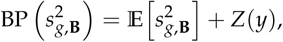

where

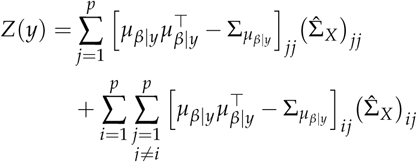

in the current population and

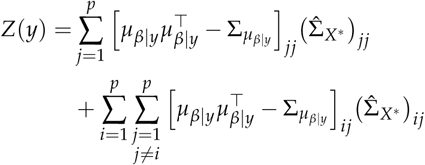

in the base population 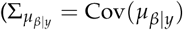.

The best predictor, therefore, consists of the unconditional expectation of the additive genomic variance (no contribution of LD) and a function that explicitly contains the weighted contribution of marker LD. This function determines whether estimators like GCTA-GREML (unconditional expectation of the random genomic variance in the base population) are biased upwards or downwards, i.e. it determines the direction and the magnitude of the bias of GCTA-GREML (this method is based on the assumption that the function *Z* constantly equals 0). In addition, we notice that this bias does not depend only on the sign of the covariance between the marker genotypes, but on the sign and the magnitude of the weighted covariances.

We emphasize that, contrary to the unconditional expectation, the best predictor maintains the structure of the additive genomic variance 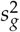 and 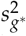, because the function *Z* can be decomposed into the weighted sample variances and covariances of the marker genotypes. Instead of the marker effects, the components of the matrix 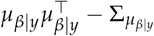 which is typically non-zero and non-diagonal, take the part of the weighting factors of the elements of 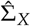 and 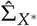. The best predictor maintains the structure of the additive genomic variance in both the current 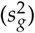 and the base population 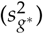 and thus conforms to the classical genetic variance (Bulmer 1971; Falconer and Mackay 1996).

The difference between the estimators *V* and *W* (*V*^*∗*^ and *W*^*∗*^, 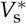 and 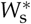) is given by the estimated *Z*(*y*) and can be obtained from Table 2. We notice that the weighted contribution of marker LD is large and positive in the case of the Arabidopsis data, whereas in the mice and wheat data the weighted contribution of marker LD is slightly negative.

To sum up, the application of the unconditional expectation of the additive genomic variance combined with the model assumptions on the marker effects in random effect models cause, at least partially, the missing contribution of LD to the estimated additive genomic variance. This goes hand in hand with the critique expressed in Kumar *et al.* (2015, 2016). It is, however, less important when estimating the additive genomic variance in the base population where the individuals are uncorrelated and less LD persists (although the marker genotypes need not be uncorrelated).

The best prediction approach eliminates the problem of the missing contribution of LD to the additive genomic variance that is caused by mathematical modeling (e.g. the assumptions in the random-effects model).

### Concluding Remarks

The variability in the genomic values and with it the additive genomic variance, is induced by the marker genotypes. The main task in investigating the random additive genomic variance in the REM is to treat the additional randomness of the genomic variance that is induced by the randomness of the marker effects. We have shown that commonly used estimators use the unconditional expectation to handle this randomness. However, we recommend the use of the best prediction approach (conditional expectation) that uses the additional information given by the genomic data, minimizes the mean square error of prediction, includes the contribution of LD, and maintains the structure of the genomic equivalent of the classical additive genetic variance.

## Acknowledgements

NS and MS were supported by the Deutsche Forschungsgemeinschaft, DFG, project number SCHL 1865/4-1, as well as the ‘RTG 1953 - Statistical Modeling of Complex Systems and Processes’. HPP was supported by the DFG project PI 377/18-1.

We would like to express our gratitude to Chris-Carolin Schön and Leo Dempfle for important remarks and advice on the paper. The remarks of anonymous reviewers helped to greatly improve the quality of this manuscript. We are also grateful to Henner Simianer, Torsten Pook and Jonas Brehmer for their comments on earlier versions of the manuscript.

## Appendix

### Genomic Best Linear Unbiased Prediction

In the REM 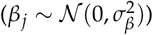 for the model

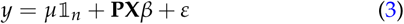

we have that

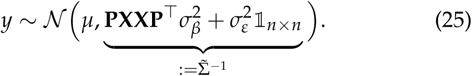

The marker effect vector *β* cannot be estimated because it is a random variable. Henderson (1984) introduced the concept of the prediction of *β*, which refers to the estimation of the realized values of the random effects. The best linear unbiased predictor (BLUP) *μ*_*β|y*_ for *β* is given by *μ*_*β|y*_ = 𝔼 [*β* | *y*] (Henderson 1984; Searle *et al.* 1992). The conditional expectation is the unique best predictor, i.e. it is unbiased and has minimal mean square error of prediction

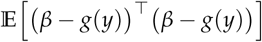

within the whole set of functions *g* that depend on the data *y* (van der Vaart 2007).

The joint distribution of *y* and *β* equals

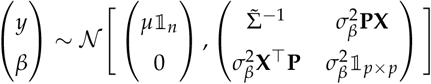

and we obtain

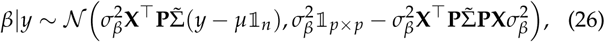

see e.g. Kotz *et al.* (2000). Consequently, the BLUP equals

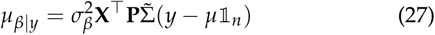

and is linear in *y*. The conditional variance-covariance matrix of *β* equals

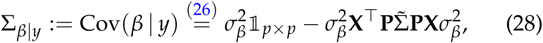

and the variance-covariance matrix of the BLUP *μ*_*β|y*_ equals

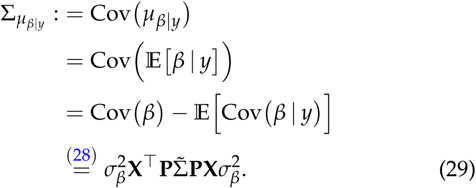

The actual estimation of the parameters in model (3) with the BLUP-method is a two-stage procedure (Das *et al.* 2004). In the first stage, a BLUE for the fixed quantities and a BLUP for the random variables are derived. However, they involve the variance components 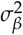 and 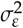 as unknown parameters. In a second stage, these parameters are replaced by estimates, and the estimators for the BLUE and the BLUP are referred to as empirical BLUE (eBLUE) and empirical BLUP (eBLUP), see Kackar and Harville (1984); Jiang (1999). Investigations on the properties of the eBLUE and the eBLUP are very complex (Searle *et al*. 1992), and often only approximate results are obtained (Kackar and Harville 1984; Jiang 1999; Das *et al*. 2004).

Assume that we are provided with estimators for the variance components using e.g. restricted maximum likelihood (REML) (Patterson and Thompson 1971; Corbeil and Searle 1976; Searle *et al*. 1992). These estimated variance components are functions of the data *y* and consequently, the eBLUE

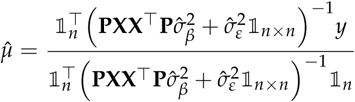

for the intercept and the eBLUP

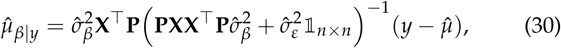

for the marker effects *β* are not even linear in the data *y* anymore (despite their naming). The unbiasedness of the estimators eBLUE and the eBLUP can be asserted if the estimated variance components 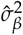 and 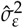 are non-negative, even functions in *y*, translation-invariant, and if the expectations of the eBLUE and eBLUP are finite (Kackar and Harville 1984). When using REML estimates for the variance components, these requirements are satisfied and the eBLUE 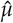 and the eBLUP 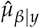 are bias-free estimators for *μ* and *β* (Jiang 1999).

Conditional on the estimation of the variance components (ignoring the randomness in the second stage of the estimation of the eBLUP), the variance-covariance matrix of the eBLUP 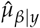 equals

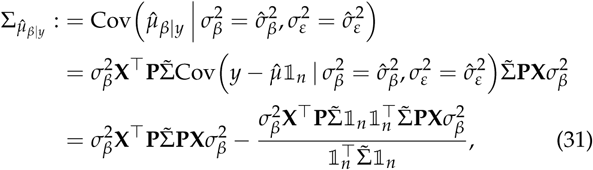

because

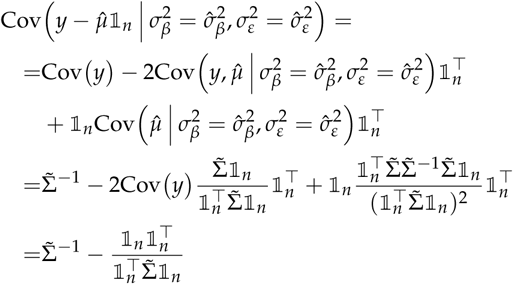

holds.

We can transfer the results derived in model (3) to model

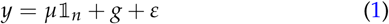

by using their equivalence in distribution. The genomic best linear unbiased predictor for *g* equals

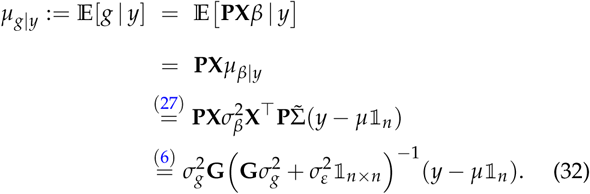

The conditional variance-covariance matrix of *g* is obtained as

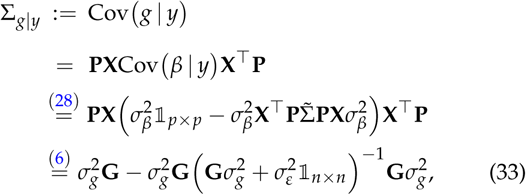

as well as the variance-covariance matrix of the BLUP *μ*_*g|y*_

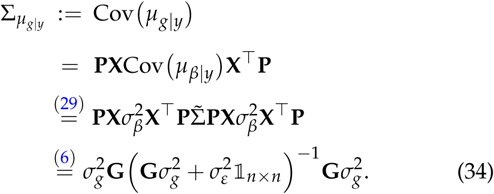

The variance-covariance matrix of the eBLUP equals

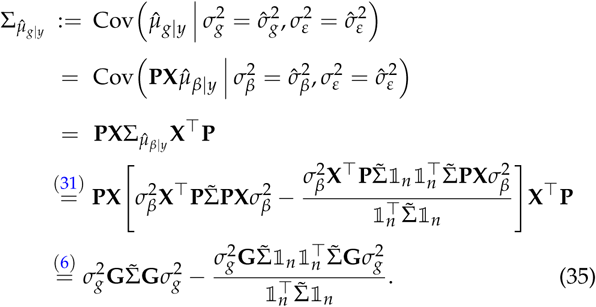

### Theoretical Variances of the Genomic Values in the REM

We review three different definitions of the theoretical variance of the genomic values in the REM (marker genotypes random, marker effects random, or both random). We focus the following analysis on the linear model (3) because of the explicit separation of marker genotypes and marker effects. For simplicity, we focus on the genomic variance in the current population. The results for the base population are obtained by replacing the data-generating process *X* with *X*^*∗*^.

#### Random Genotypes and Random Effects

If the marker genotypes as well as the marker effects are the source of genomic variation, we calculate the variance of the genomic value according to the law of total variance as:

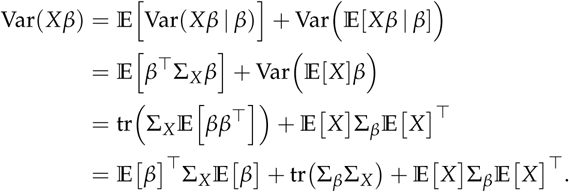

The unconditional expectation and the variance operator in the second line apply to the random marker effect vector *β*. Because of the model assumptions on the marker effects in (5) and the mean-centered marker genotypes (𝔼 [*X*] = 0), we obtain

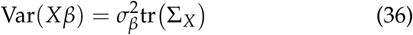

with the interpretation as the variance of a randomly sampled (representative) individual for a trait with random effects.

#### Fixed Genotypes and Random Effects

If the genomic variation is caused by the marker effects only and the marker genotypes are fixed, then the *n*-vector of genomic values is normally distributed:

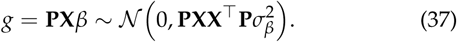

In order to obtain an average theoretical variance of the individuals in the sample, we calculate the mean trace of the variance-covariance matrix of the genomic values:

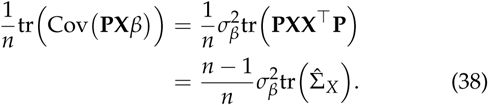

This approximately equals the variance of a randomly sampled individual for a randomly sampled trait, see (36). Even when the marker genotypes are fixed, their sample variance-covariance matrix contributes to the theoretical variance of the genomic values when averaging over the individuals in the sample.

#### Random Genotypes and Fixed Effects

Probably the most common assumption on the nature of the genome is that the marker genotypes are random, whereas the marker effects are fixed (Falconer and Mackay 1996). In order to translate these assumption to the variance of the genomic values in the REM, we have to condition on the marker effects (i.e. we fix them). Then, the theoretical variance of the genomic values of a individual with random marker genotypes (representative individual) and with fixed marker effects equals

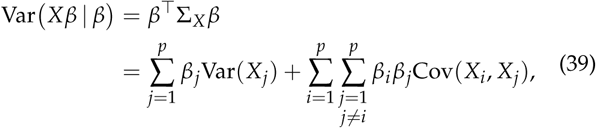

and describes the genomic equivalent of the definition of the additive genetic variance (Bulmer 1971; Falconer and Mackay 1996).

### Estimation of the Additive Genomic Variance in the REM

In sections The Expectation of the Additive Genomic Variance and Best Prediction of the Additive Genomic Variance we have introduced ways to predict the random additive genomic variance

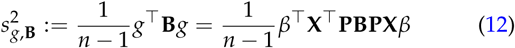

in the REM, namely by using the unconditional expectation

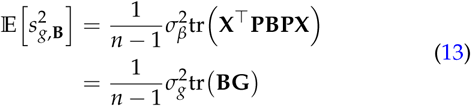

and the best predictor

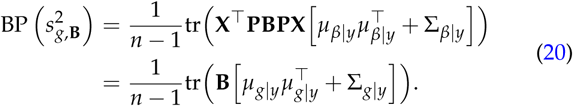

In the following, we introduce estimators for these quantities and investigate their properties.

#### Estimation of the Unconditional Expectation

For any given positive semi-definite matrix **B**, the unconditional expectation

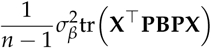

in model (3) can be estimated by

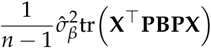

after having obtained an estimate 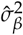 of the variance component 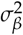. In the equivalent model (1), we can estimate

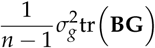

by using

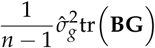

for any positive semi-definite matrix **B**.

The specification of **B** as in Section The Expectation of the Additive Genomic Variance leads to the explicit form of the estimators

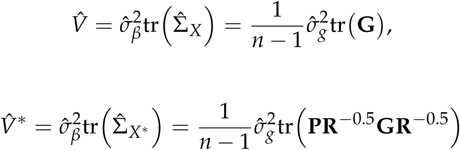

and

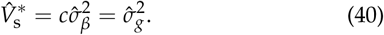

#### Empirical Best Prediction (eBP)

Because of equalities (28) and (29) and the variance-covariance matrix 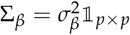 of *β* we have that

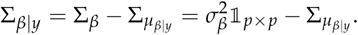

Consequently, the best predictor of 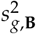 defined in (20) equals

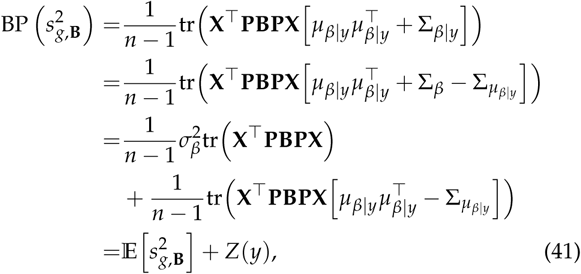

where

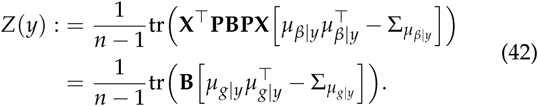

We have partitioned the best predictor of 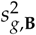 into the unconditional expectation of 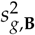 and the random variable *Z* which is realized in the phenotypic data *y*. The random variable *Z* specifies the adaption of the best predictor to the data and incorporates the contribution of (marker) LD. The expectation of *Z* over all possible data *y* is 0 because

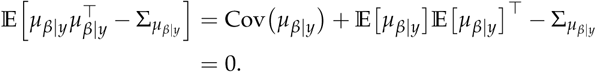

The sign of the realization of *Z* determines whether the best predictor is larger (positive weighted LD) or smaller (negative weighted LD) than the unconditional expectation.

The task of finding an eBP for 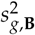 is reduced to estimating the realized values of *Z* because of the connection derived in (41). We replace the BLUP and their variance-covariance matrix in equation (42) by the eBLUP and its estimated variance-covariance matrix:

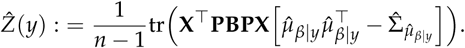

We assume that we are provided with REML-estimators 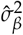 and 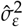 for the variance components. Then, we find

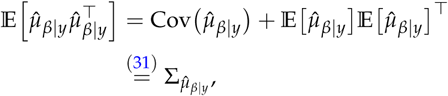

because the eBLUP is unbiased for 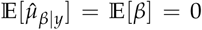 (Jiang 1999). Unfortunately, the unbiasedness of the estimated variance-covariance matrix of the eBLUP can only be asserted in a trivial way by conditioning on the estimated variance components:

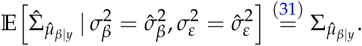

Therefore, the expectation of 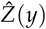 (conditionally on the variance components) equals 0.

The same holds true for

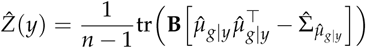

in the equivalent model, because the quantities 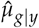 and 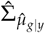 are linear combinations of 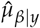 and 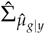, see (32) and (35).

Altogether, we can define the unbiased (conditionally on the estimated variance components) empirical best predictor

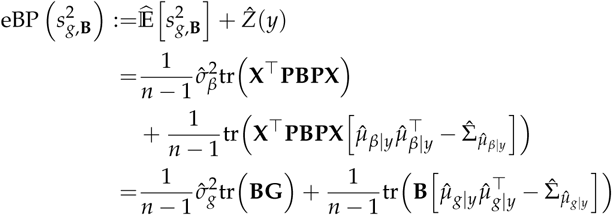

for the additive genomic variance 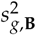.

The specification of **B** as in Section Best Prediction of the Additive Genomic Variance leads to the explicit form

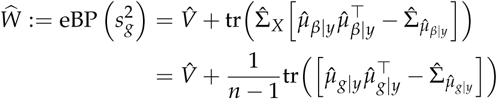

of the eBP for the additive genomic variance in the current population, and to the eBP

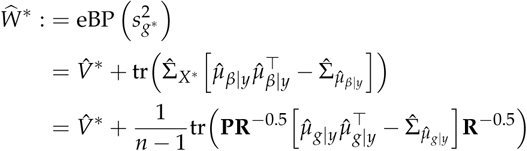

for the additive genomic variance in the base population.

Using the GRM **G** for a transformation to the base population is not well-defined because **G** is singular. However, because 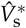, see (40), is commonly used, we want to find an analogous formula for the empirical best predictor in this set-up.

Instead of calculating

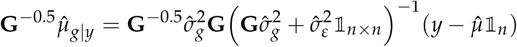

and

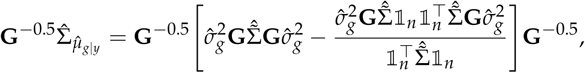

we use

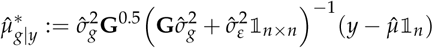

and

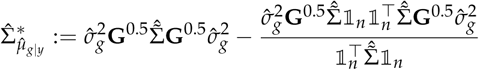

as substitutes. Then, we define

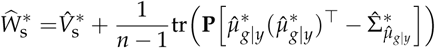

as an approximation of the empirical best predictor of the additive genomic variance in the base population when using the GRM for the transformation.

