## Supplement 1 for "Best Prediction of the Additive Genomic Variance in Random-Effects Models"

### Supporting Material for: "Best Prediction of the Additive Genomic Variance in Random-Effects Models"

28. Februar 2019

#### 1 Estimation of the Unconditional Expectation and the Best Predictor of the Additive Genomic Variance in the "Equivalent Linear Model"

We review the formulae for the estimated unconditional expectation as well as the empirical best predictor of the additive genomic variance introduced in the paper, and provide code for an implementation using the free software *R*.

We focus on the linear model

$$y = \mu \mathbb{1}_n + g + \varepsilon,$$

for the phenotypic values $y$ where $\mu$ denotes a fixed intercept, $\mathbb{1}_n := (1, ..., 1)^\top$ is an $n$-row-vector, $\varepsilon \sim \mathcal{N}(0, \sigma_\varepsilon^2 \mathbb{1}_{n\times n})$, and $\mathbb{1}_{n\times n}$ is the $n$-dimensional identity matrix.
In the random-effects model, the mean-centered ($\bar{g} := \frac{1}{n}\mathbb{1}_n^\top g = 0$) genomic values $g$ are distributed as

$$g \sim \mathcal{N}\big(0, \sigma_g^2 \mathbf{G}\big).$$

When using genomic marker data, the genomic relationship matrix $\mathbf{G}$ can be defined as

$$\mathbf{G} := \mathbf{P}\mathbf{X}\mathbf{X}^\top \mathbf{P}/c \tag{1}$$

where $c > 0$ is a constant, $\mathbf{X}$ the marker genotypes matrix and

$$\mathbf{P} := \mathbb{1}_{n\times n} - \mathbb{1}_n \mathbb{1}_n^\top / n \tag{2}$$

carries out the column-wise mean-centering of the marker genotypes. Let $\sigma_\beta^2$ be the variance component of the marker effects $\beta$. Then, the variance component in the equivalent model equals $\sigma_g^2 = c\sigma_\beta^2$ and the constant $c$ is usually defined as

$$c := 2 \sum p_j (1 - p_j), \tag{3}$$

where $p_j$ is the frequency of the minor allele at marker $j$. We use the sample variance of the phenotypic values

$$\hat{\sigma}_y^2 := \frac{1}{n-1} y^\top \mathbf{P} y \tag{4}$$

to scale the phenotypic values to unit variance by setting

$$\tilde{y} = \frac{y}{\hat{\sigma}_y}. \tag{5}$$

We fit the model using the $R$-package "sommer" which returns the estimated variance components $\hat{\sigma}_g^2$ and $\hat{\sigma}_\varepsilon^2$ (REML approach). In addition, we obtain the empirical BLUP (eBLUP)

$$\hat{\mu}_{g|y} = \hat{\sigma}_g^2 \mathbf{G} \big( \mathbf{G} \hat{\sigma}_g^2 + \hat{\sigma}_\varepsilon^2 \mathbb{1}_{n \times n} \big)^{-1} (y - \hat{\mu} \mathbb{1}_n) \tag{6}$$

for the random vector $g$ and the estimated variance-covariance matrix

$$\begin{aligned} \hat{\Sigma}_{\hat{\mu}_{g|y}} =& \hat{\sigma}_g^2 \mathbf{G} \big( \mathbf{G} \hat{\sigma}_g^2 + \hat{\sigma}_\varepsilon^2 \mathbb{1}_{n \times n} \big)^{-1} \mathbf{G} \hat{\sigma}_g^2 \\ &- \frac{\hat{\sigma}_g^2 \mathbf{G} \big( \mathbf{G} \hat{\sigma}_g^2 + \hat{\sigma}_\varepsilon^2 \mathbb{1}_{n \times n} \big)^{-1} \mathbb{1}_n \mathbb{1}_n^\top \big( \mathbf{G} \hat{\sigma}_g^2 + \hat{\sigma}_\varepsilon^2 \mathbb{1}_{n \times n} \big)^{-1} \mathbf{G} \hat{\sigma}_g^2}{\mathbb{1}_n^\top \big( \mathbf{G} \hat{\sigma}_g^2 + \hat{\sigma}_\varepsilon^2 \mathbb{1}_{n \times n} \big)^{-1} \mathbb{1}_n} \end{aligned} \tag{7}$$

of the eBLUP $\hat{\mu}_{g|y}$ of $g$.

When interest is in the current population, we have to set

$$\mathbf{B}_{\mathrm{c}} := \mathbb{1}_{n \times n},$$

and obtain the estimated unconditional expectation

$$\hat{V} = \frac{1}{n-1} \hat{\sigma}_g^2 \operatorname{tr}(\mathbf{G}) \tag{8}$$

and the empirical best predictor (eBP)

$$\widehat{W} = \hat{V} + \frac{1}{n-1} \hat{\mu}_{g|y}^\top \hat{\mu}_{g|y} - \frac{1}{n-1} \operatorname{tr}\Big( \hat{\Sigma}_{\hat{\mu}_{g|y}} \Big) \tag{9}$$

for the additive genomic variance in the current population.

We define estimators for the heritabilities in the current population by dividing the estimated unconditional expectation

$$\hat{h}_V^2 := \frac{\hat{V}}{\hat{\sigma}_y^2} = \hat{V} \tag{10}$$

as well as the eBP

$$\hat{h}_W^2 := \frac{\widehat{W}}{\hat{\sigma}_y^2} = \widehat{W}. \tag{11}$$

of the additive genomic variance by the phenotypic sample variance.
Alternatively, we can assume that the phenotypic variance equals the sum of the estimated genomic samples variances in the current population and the estimated residual variance:

$$\tilde{\sigma}_{y,V}^2 := \hat{V} + \hat{\sigma}_\varepsilon^2 \tag{12}$$

and

$$\tilde{\sigma}_{y,W}^2 := \widehat{W} + \hat{\sigma}_\varepsilon^2. \tag{13}$$

These estimators for the phenotypic variance depend on the specific choice of the estimator for the genomic sample variance and lead to alternative heritability estimates:

$$\tilde{h}_V^2 := \frac{\hat{V}}{\tilde{\sigma}_{y,V}^2} \tag{14}$$

and

$$\tilde{h}^2_W := \frac{\widehat{W}}{\tilde{\sigma}^2_{y,W}}. \tag{15}$$

When interest is in the base population, we have to set

$$\mathbf{B}_{\mathrm{b}} := \mathbf{R}^{-0.5}\mathbf{P}\mathbf{R}^{-0.5}, \tag{16}$$

and we obtain the estimated unconditional expectation

$$\hat{V}^* = \frac{1}{n-1}\hat{\sigma}^2_g \mathrm{tr}\Big(\mathbf{R}^{-0.5}\mathbf{P}\mathbf{R}^{-0.5}\mathbf{G}\Big) \tag{17}$$

and the eBP

$$\widehat{W}^* = \hat{V}^* + \frac{1}{n-1}\hat{\mu}^\top_{g|y}\mathbf{R}^{-0.5}\mathbf{P}\mathbf{R}^{-0.5}\hat{\mu}_{g|y} - \frac{1}{n-1}\mathrm{tr}\Big(\mathbf{R}^{-0.5}\mathbf{P}\mathbf{R}^{-0.5}\hat{\Sigma}_{\hat{\mu}_{g|y}}\Big). \tag{18}$$

for the additive genomic variance in the base population.

The GRM $\mathbf{G}$ is singular and therefore $\mathbf{G}^{-0.5}$ is not well-defined. However, it is possible to decompose $\mathbf{G}$ into

$$\mathbf{G} = \mathbf{G}^{0.5}\mathbf{G}^{0.5}.$$

Then, we directly set the unconditional expectation of the additive genomic variance to

$$\hat{V}^*_{\mathrm{s}} = \hat{\sigma}^2_g \tag{19}$$

instead of calculating

$$\frac{1}{n-1}\hat{\sigma}^2_g \mathrm{tr}\Big(\mathbf{P}\mathbf{G}^{-0.5}\mathbf{G}\mathbf{G}^{-0.5}\Big) = \frac{1}{n-1}\hat{\sigma}^2_g \mathrm{tr}(\mathbf{P}) = \hat{\sigma}^2_g$$

when using the GRM for the transformation to the base population in (17).
We proceed analogously for the eBP. Instead of calculating

$$\mathbf{G}^{-0.5}\hat{\mu}_{g|y} = \mathbf{G}^{-0.5}\hat{\sigma}^2_g\mathbf{G}\Big(\mathbf{G}\hat{\sigma}^2_g + \hat{\sigma}^2_\varepsilon \mathbb{1}_{n\times n}\Big)^{-1}(y - \hat{\mu}\mathbb{1}_n)$$

and

$$\mathbf{G}^{-0.5}\hat{\Sigma}_{\hat{\mu}_{g|y}} = \mathbf{G}^{-0.5}\left[\hat{\sigma}^2_g\mathbf{G}\hat{\hat{\Sigma}}\mathbf{G}\hat{\sigma}^2_g - \frac{\hat{\sigma}^2_g\mathbf{G}\hat{\hat{\Sigma}}\mathbb{1}_n\mathbb{1}_n^\top\hat{\hat{\Sigma}}\mathbf{G}\hat{\sigma}^2_g}{\mathbb{1}_n^\top\hat{\hat{\Sigma}}\mathbb{1}_n}\right]\mathbf{G}^{-0.5}$$

in (18), we use

$$\hat{\mu}^*_{g|y} := \hat{\sigma}^2_g\mathbf{G}^{0.5}\Big(\mathbf{G}\hat{\sigma}^2_g + \hat{\sigma}^2_\varepsilon \mathbb{1}_{n\times n}\Big)^{-1}(y - \hat{\mu}\mathbb{1}_n) \tag{20}$$

and

$$\hat{\Sigma}^*_{\hat{\mu}_{g|y}} := \hat{\sigma}^2_g\mathbf{G}^{0.5}\hat{\hat{\Sigma}}\mathbf{G}^{0.5}\hat{\sigma}^2_g - \frac{\hat{\sigma}^2_g\mathbf{G}^{0.5}\hat{\hat{\Sigma}}\mathbb{1}_n\mathbb{1}_n^\top\hat{\hat{\Sigma}}\mathbf{G}^{0.5}\hat{\sigma}^2_g}{\mathbb{1}_n^\top\hat{\hat{\Sigma}}\mathbb{1}_n} \tag{21}$$

as surrogates. Then, we define

$$\widehat{W}^*_{\mathrm{s}} = \hat{V}^*_{\mathrm{s}} + \frac{1}{n-1}\big(\hat{\mu}^*_{g|y}\big)^\top\mathbf{P}\hat{\mu}^*_{g|y} - \frac{1}{n-1}\mathrm{tr}\Big(\mathbf{P}\hat{\Sigma}^*_{\hat{\mu}_{g|y}}\Big) \tag{22}$$

for the eBP of the additive genomic variance in the base population when using the GRM for the transformation.

#### 2 R-Script for the Estimation of the Additive Genomic Variance

We implement the quantities defined above using the following $R$-code.

```
################################################################################
### Estimation of the Genomic Variance in the ‘‘Equivalent Linear Model’’ ######
################################################################################

# Author
# Nicholas Schreck,

# Estimation of the Genomic Variance in the "Equivalent Linear Model"

# Copyright (C) 2018 -- 2019 Nicholas Schreck

# This program is free software; you can redistribute it and/or
# modify it under the terms of the GNU General Public License
# as published by the Free Software Foundation; either version 2
# of the License, or (at your option) any later version.

# This program is distributed in the hope that it will be useful,
# but WITHOUT ANY WARRANTY; without even the implied warranty of
# MERCHANTABILITY or FITNESS FOR A PARTICULAR PURPOSE.  See the
# GNU General Public License for more details.

# You should have received a copy of the GNU General Public License
# along with this program; if not, write to the Free Software
# Foundation, Inc., 59 Temple Place - Suix2te 330, Boston, MA  02111-1307, USA.

################################################################################
# Packages
library(sommer)
library(BGLR)
################################################################################
# Genomic data (phenotypes and marker genotypes)
data(mice)
data(wheat)
load("Arabidopsis1001genomes_1057lines_193697SNPs.Rdata")
y.all <- list(mice.pheno$Obesity.BodyLength, wheat.Y[, 1],
              phenogeno$pheno$FT10_mean) # Phenotypic values
Ped.all <- list(mice.A, wheat.A, NA) # Pedigree matrices
X.all <- list(mice.X, wheat.X, phenogeno$geno) # marker-genotyp matrices
nr.datasets <- length(y.all)
################################################################################
# Output Variables
Variables <- c("eps", "V_hat", "h2_V", "sum_V", "h2_V_sum", "W_hat", "h2_W",
               "sum_W", "h2_Wsum", "Vstar_hat", "Wstar_hat", "Vstars_hat",
               "Wstars_hat")
nr.variables <- length(Variables)
result <- matrix(NA, nr.variables, nr.datasets)
rownames(result) <- Variables
colnames(result)  <- c("Mice", "Wheat", "Arabidopsis")
################################################################################
rm(mice.A, mice.pheno, mice.X, wheat.A, wheat.X, wheat.Y, phenogeno, wheat.sets,
   Variables, nr.variables)

################################################################################
```

```r
###############################################################################
# Model fit and estimation of genomic variances
for (j in 1 : nr.datasets){ # for each dataset
  # Preliminaries
  y <- y.all[[j]] # fix phenotype
  y.scaled <- y / sd(y) # y scaled to variance of 1, see equation (5)
  n <- length(y)
  ones <- matrix(1, nrow=n, ncol=1)
  P <- diag(1, n, n) - matrix(1, n, n) / n # Equation (2)
  X <- X.all[[j]] # fix marker-genotypes
  p <- dim(X)[2]
  c <- sum( colMeans(X) * (1 - colMeans(X) / 2) ) # vanRaden c, see equation (3)
  G <- P %*% tcrossprod(X) %*% P / c # equation (1)
  rm(X)
  #############################################################################
  # Fit gBLUP model
  lm.equi <- mmer(Y=y.scaled, X=ones, Z=list(A=list(K=G)), silent=TRUE,
                  iters=50, tolpar=1e-7, tolparinv=1e-9, date.warning = FALSE)
  var.g.hat <- as.numeric(lm.equi$var.comp$A) # variance component g
  var.e.hat <- as.numeric(lm.equi$var.comp$units) # variance comp. eps
  mu.hat <- rep(lm.equi$beta.hat, n) # Estimate of Intercept
  g.hat <- as.vector(lm.equi$u.hat$A)  # eBLUP, equation (6)
  cov.g.hat <- as.matrix(lm.equi$Var.u.hat$A$T1) # Cov of eBLUP, equation (7)
  rm(lm.equi)
  # Genomic variances in the current population
  gv_cur <- GenVarCur(GRM=G, varg=var.g.hat, eBLUP=g.hat, eBLUPcov=cov.g.hat)
  V.hat <- gv_cur[1]
  h2.V <- V.hat # formula (10)
  sum.V <- V.hat + var.e.hat # formula (12)
  h2.V.sum <- V.hat / sum.V # formula (14)
  W.hat <- gv_cur[2]
  h2.W <- W.hat # (11)
  sum.W <- W.hat + var.e.hat # formula (13)
  h2.W.sum <- W.hat / sum.W # formula (15)
  # Genomic variances in the base population (via Pedigree)
  Ped <- Ped.all[[j]]
  if(!is.na(Ped)){
    gv_base_ped <- GenVarBase_Ped(PedMat=Ped, GRM=G, varg=var.g.hat, eBLUP=g.hat,
                                  eBLUPcov=cov.g.hat)
    V.hat.star <- gv_base_ped[1]
    W.hat.star <- gv_base_ped[2]
  }else{
     V.hat.star <- W.hat.star <- NA
  }
  # Genomic variances in the base population (via GRM)
  gv_base_grm <- GenVarBase_GRM(GRM=G, phenos=y.scaled, mu=mu.hat,
                                varg=var.g.hat, eBLUP=g.hat, eBLUPcov=cov.g.hat,
                                vare=var.e.hat)
  V.hat.star.s <- gv_base_grm[1]
  W.hat.star.s <- gv_base_grm[2]
  ###############################################################################
  result[, j] <- c(var.e.hat, V.hat, h2.V, sum.V, h2.V.sum, W.hat, h2.W,
                  sum.W, h2.W.sum, V.hat.star, W.hat.star, V.hat.star.s,
                  W.hat.star.s)
  rm(var.e.hat, V.hat, h2.V, sum.V, h2.V.sum, W.hat, h2.W, sum.W, h2.W.sum,
     V.hat.star, W.hat.star, V.hat.star.s, W.hat.star.s, y, y.scaled, var.g.hat,
     g.hat, cov.g.hat)
```

```r
}
################################################################################
################################################################################
# Functions for the calculation of the genomic variance in different set-ups
GenVarCur <- function(GRM, varg, eBLUP, eBLUPcov){
  n <- dim(GRM)[[1]]
  V <- varg * sum( diag(GRM) ) / (n - 1) # formula (8)
  W <- V +  sum(eBLUP ^ 2) / (n - 1) - sum( diag(eBLUPcov) ) / (n - 1)
  return(c(V, W))
}
GenVarBase_Ped <- function(PedMat, GRM, varg, eBLUP, eBLUPcov){
  n <- dim(GRM)[[1]]
  P <- diag(1, n, n) - matrix(1, n, n) / n # Equation (2)
  E <- eigen(PedMat)
  Q <- E$vectors
  D <- E$values
  Ped.inv.halve <- Q %*% diag( 1 / sqrt(D), nrow(Q), nrow(Q)) %*% t(Q)
  B_b <- Ped.inv.halve %*% P %*% Ped.inv.halve # equation (16)
  V <- varg * sum( diag(B_b %*% GRM) ) / (n - 1) # formula (17)
  W <- V + t(eBLUP) %*% B_b %*% eBLUP /
    (n - 1) - sum(diag(B_b %*% eBLUPcov)) / (n - 1) # formula (18)
  return(c(V, W))
}
GenVarBase_GRM <- function(GRM, phenos, mu, varg, eBLUP, eBLUPcov, vare){
  n <- dim(GRM)[[1]]
  P <- diag(1, n, n) - matrix(1, n, n) / n # Equation (2)
  E_G <- eigen(GRM)
  Q_G <- E_G$vectors
  D_G <- E_G$values
  D_G[D_G < 0] <- 0 # Last EV might be -0
  G_halve <- Q_G %*% diag(sqrt(D_G), nrow(Q_G), nrow(Q_G)) %*% t(Q_G)
  Cov.y.minus <- chol2inv(GRM * varg + diag(vare, n, n))
  eBLUP.s <- varg * G_halve %*% Cov.y.minus %*% (phenos - mu)
  # formula (20)
  eBLUPcov.s <- varg ^ 2 * G_halve %*% Cov.y.minus %*% G_halve -
    (varg ^ 2 / sum(Cov.y.minus)) * G_halve %*% Cov.y.minus %*%
    matrix(1, n, n) %*% Cov.y.minus %*% G_halve # formula (21)
  V <- varg # formula (19)
  W <- V + t(eBLUP.s) %*% P %*% eBLUP.s / (n - 1) - sum(diag(P %*%
        eBLUPcov.s)) / (n - 1) # formula (22)
  return(c(V, W))
}
```

#### 3 Results

We present the outcome of the *R*-script above in table format.

**Table S1**: Estimation results for the unconditional expectation $V$ and the best predictor $W$ for the additive genomic variance $s_g^2$ in the current population for the mice, wheat, and Arabidopsis datasets. We also present the corresponding heritabilities with respect to the sample variance of the phenotypic values and with respect to the sum of the additive genomic and residual variance $\sigma_\varepsilon^2$.
In addition, we depict the estimation results for the unconditional expectation $V^*$ ($V_s^*$ when using the GRM for the transformation) and the best predictor $W^*$ ($W_s^*$ when using the GRM for the transformation) for the additive genomic variance in the base population.

| Genom. Var. / Heritab. \ Data | Population | Mice | Wheat | Arabidopsis |
|---|---|---|---|---|
| $\hat{V}\ (=\hat{h}_V^2)$ | Current | 0.3737749 | 0.6039708 | 0.47333803 |
| $\hat{V}+\hat{\sigma}_\varepsilon^2$ | | 1.0754963 | 1.1449704 | 0.54832029 |
| $\tilde{h}_V^2$ | | 0.3475371 | 0.5274990 | 0.86325098 |
| $\widehat{W}\ (=\hat{h}_W^2)$ | | 0.2982787 | 0.4590001 | 0.92501779 |
| $\widehat{W}+\hat{\sigma}_\varepsilon^2$ | | 1.0000002 | 0.9999998 | 1.00000005 |
| $\tilde{h}_W^2$ | | 0.2982787 | 0.4590002 | 0.92501774 |
| $\hat{V}^*$ | Base | 0.3704021 | 3.0621134 | – |
| $\widehat{W}^*$ | | 0.3089758 | 2.0095836 | – |
| $\hat{V}_s^*$ | | 0.3639248 | 1.3158006 | 0.80762011 |
| $\widehat{W}_s^*$ | | 0.3577692 | 1.2300300 | 1.30240520 |
